# Microbial community composition interacts with local abiotic conditions to drive colonization resistance in human gut microbiome samples

**DOI:** 10.1101/2020.08.18.255695

**Authors:** Michael Baumgartner, Katia R Pfrunder-Cardozo, Alex R Hall

## Abstract

Biological invasions can alter ecosystem stability and function, and predicting what happens when a new species or strain arrives remains a major challenge in ecology. In the mammalian gastrointestinal tract, susceptibility of the resident microbial community to invasion by pathogens has important implications for host health. However, at the community level, it is unclear whether susceptibility to invasion depends mostly on resident community composition (which microbes are present), or also on local abiotic conditions (such as nutrient status). Here, we used a gut microcosm system to disentangle some of the drivers of susceptibility to invasion in microbial communities sampled from humans. We found resident microbial communities inhibited an invading *E. coli* strain, compared to community-free control treatments, sometimes excluding the invader completely (colonization resistance). These effects were stronger at later time points, coinciding with shifts in microbial community composition and nutrient availability. By separating these two components (microbial community and abiotic environment), we found taxonomic composition played a crucial role in suppressing invasion, but this depended critically on local abiotic conditions (adapted communities were more suppressive in nutrient-depleted conditions). This helps predict when resident communities will be most susceptible to invasion, with implications for optimizing treatments based around microbiota management.

## Introduction

Biological invasions have major impacts on ecosystem function and diversity [1]. Many factors can influence whether invading species successfully colonize new communities [2,3], making it a key challenge in ecology to understand what drives the outcome of invasions. One ecosystem where susceptibility to invasion has direct impacts on human health is the mammalian intestinal microbiome. Here, the resident microbial community protects against infection by pathogens, and disturbance can lead to opportunistic invasion / infection [4–7]. What determines the ability of resident gastrointestinal communities to suppress colonization by invading, non-resident species (colonization resistance [8,9])? This question is central to our basic understanding of how microbial communities respond to invasions. It is also important for explaining variable susceptibility to infectious disease, and predicting the success of microbiota-based therapies such as faecal microbial transplantation [10,11]. In some cases, physiological mechanisms by which individual resident taxa impact colonization resistance have been characterized [12], including direct interactions among microbial species and indirectly via interplay with the host immune system [13]. Other recent studies have demonstrated the net effect of entire microbial communities on colonization success of invading strains [14,15]. However, it is unclear whether such effects depend solely on which resident organisms are present (e.g. particular strains or species), in which case they could potentially be predicted from metagenomic data, or also on local abiotic conditions, such as nutrient availability. Disentangling these factors is challenging, because in natural microbiomes they are intertwined: changing community structure modifies the local micro-environment, and vice versa.

Depending on which of the known types of microbial interactions involved in colonization resistance are most important, we can expect susceptibility of resident microbiota to invasion to depend on microbial community composition and/or local abiotic conditions in various ways. For example, if invading strains are inhibited by toxins produced by a subset of the resident bacteria, then colonization resistance will rely primarily on the presence of particular taxa (those encoding invader-inhibiting mechanisms). There is support for such mechanisms in that the type VI secretion system, encoding contact-dependent growth inhibition of other strains, is widespread in the common gut phylum Bacteroidetes [16–18]. Similarly, production of narrow-spectrum antibacterial toxins is common in the human gut microbiome, and gut commensals producing bacteriocins, such as *Escherichia coli* and *Bifidobacterium* spp., can suppress pathogens compared to non-producing mutant strains [19,20]. By contrast, other mechanisms of colonization resistance should be less contingent on community composition. For example, resource competition between resident microbiota and invading strains does not necessarily require a specific set of taxa to be present, only that the resident community is diverse and dense enough to scavenge shared resources effectively. In support of a role for resource competition, after disturbance of microbiota by antibiotic treatment the availability of free sugars and amino acids can increase, and subsequently be exploited by opportunistic pathogens like *Salmonella enterica* serovar Typhimurium or *Clostridium difficile* [21,22]. Crucially, we can expect this type of colonization resistance to depend strongly on local abiotic conditions, in that resource competition will be most effective at inhibiting invading strains when resources become scarce [23].

We aimed to test whether colonization resistance in individual human microbiome samples is affected by changes over time in microbiota composition, local abiotic conditions (in particular, nutrient limitation), or both. We did this by observing invasion of human-associated microbiota by a non-resident focal strain in replicated microcosm experiments. We used a gut microcosm system [14] to co-cultivate the invading focal strain with microbiota sampled from three healthy human donors. We used *E. coli* as the focal strain because it is a common gut commensal [24], but also an opportunistic pathogen [25,26]. By using microcosms with sterilized and ‘live’ versions of the resident microbiota, we quantified the effect of resident microbiota on growth of our focal strain (estimated by selective plating). We maintained each microcosm with periodic sampling for 72h. This allowed us to monitor the effect of resident microbiota both early on and during later stages when we expected the environment to have become relatively nutrient-poor. We also tracked changes in community composition over time, using amplicon sequencing. Finally, by separating resident microorganisms from the supernatant (liquid phase) of individual microcosms and then using these two phases in further experiments, we were able to disentangle the effects of the resident microbiota and the local abiotic conditions on growth of the focal strain. Our results show that the microbiomes of three human donors successfully suppressed our focal strain, in some cases amounting to full colonization resistance (exclusion of the invading strain), but this suppressive effect was contingent on changes in both resident microbial community composition and changing environmental conditions over time.

## Materials and methods

### Focal bacterial strain (invader)

We used *Escherichia coli* K-12 MG1655 with a streptomycin resistance mutation (*rpsL* K43R) as our focal strain. Prior to the main microcosm experiment, we inoculated the focal strain for 24h in anaerobic basal medium [27,28] with some modifications (2 g/l Peptone, 2 g/l Tryptone, 2 g/l Yeast extract, 0.1 g/l NaCl, 0.04g K_2_HPO_4_, 0.04 g/l KH_2_PO_4_, 0.01 g/l MgSO_4_x7H_2_O, 0.01 g/l CaClx6H_2_O, 2g/l NaHCO_3_, 2 ml Tween 80, 0.005 g/l Hemin, 0.5 g/l L-Cysteine, 0.5 g/l bile salts, 2g/l Starch, 1.5 g/l casein, 0.001g/l Resazurin, pH adjusted to 7, addition of 0.001g/l Menadion after autoclaving) under a constant stream of nitrogen gas, at 37 °C and 220 rpm in a shaking incubator.

### Faecal samples

The following protocol for obtaining human faecal samples was approved by the ETH Zürich Ethics Commission (EK 2016-N-55). We collected samples on 2018-01-08 from three anonymous, consenting Donors and kept them anaerobic for approximately one hour until sampling was finished. We then re-suspended 10 g of each sample in 100 ml anaerobic peptone wash (1 g/l peptone, 0.5 g/l L-Cysteine, 0.5 g/l bile salts, 0.001 g/l Resazurin) by stirring for 5 minutes followed by 15 min of resting to sediment. 50 ml of each of these 10% (w/v) faecal slurries were transferred to 100 ml flasks and autoclaved to prepare sterile slurry. The other 50 ml, with ‘live’ slurry, were stored at room temperature until further processing, similar to the procedure described in [14].

### Anaerobic batch culture system, sampling and bacterial enumeration

Microcosms consisted of Hungate tubes, as described in [14]. We filled each tube with 7.2 ml basal medium, flushed the head space with nitrogen gas, and then autoclaved. We then added either 850 µl of sterilized slurry (community-free treatments) or 350 µl of live slurry and 500 µl of sterilized slurry (community treatments). We also added the focal strain to each microcosm, by adding 8 µl of overnight culture (approximately 10^6^ CFU ml^-1^). For the control treatment (basal medium without faecal slurry), we inoculated the focal strain in basal medium supplemented with 850 µl of peptone wash, to equalize the volume with the human donor treatments. We incubated all microcosms at 37 °C in a static incubator and took samples after 2, 24, 48 and 72 h. To estimate focal strain abundance, we serially diluted samples in phosphate-buffered solution (PBS) and plated on Chromatic MH agar (Liofilchem, Roseto, Italy) supplemented with Streptomycin (100 µg/ml), before counting colony forming units. We initially screened the faecal slurry of each human donor to verify the specificity of our selective plates; this revealed no resident bacteria able to grow on these plates. To estimate total bacterial abundance (including the resident microbiota) we used flow cytometry. We diluted each microcosm sample with PBS and stained it with Sybr green (Life Technologies, Zug, Switzerland), with a final concentration of 10^−4^ of the commercial stock solution. We used a Novocyte 2000R (ACEA Biosciences Inc., San Diego, USA) equipped with a laser emitting at 488 nm and the standard filter setup to measure cell density. Detection of bacteria was based on their signature in a plot of forward scatter versus green fluorescence.

### Supernatant experiment

We extracted the supernatant of each microcosm at the end of the experiment, from both community- and community-free treatments. As a control treatment, we supplemented fresh basal medium with thawed, sterilized slurry from each human donor of the batch culture experiment, the same way as for microcosms at the start of the main experiment in the community-free treatments. To extract supernatants from each treatment (community, community-free and control), we transferred 1.5 ml of each culture to a 2 ml tube and centrifuged at 10,000 rpm for 5 min. Each supernatant was then transferred to a syringe and sterile filtered (0.22 µm pore size) into a fresh tube. We then transferred 250 µl aliquots of each sterile-filtered supernatant into a 96-well microplate and inoculated each well with 5 µl of an overnight culture of our focal strain. We incubated the microplate for 24h at 37 °C without shaking. We took samples for CFU counts at the beginning and end of the experiment, and when necessary diluted them in PBS before plating on LB plates. Note this inhibition assay was aerobic, whereas the main experiment and the below experiment with adapted/fresh communities were anaerobic.

### Colonization resistance of adapted and fresh communities

To test whether resident microbial communities that had adapted to the microcosm environment were more resistant against invasion by our focal strain, and whether this effect was contingent on changes in the abiotic conditions in each microcosm over time, we produced various combinations of communities and supernatants. First, we produced and then froze down community samples that were either adapted (had been incubated for 72h in the absence of the focal strain, but in the same conditions as in the main microcosm experiment above) or fresh (prepared as at the start of the community treatments above, but without the focal strain). Using frozen community samples here allowed us to directly compare samples from before and after 72h cultivation. There is a risk some taxa are affected by freeze-thawing, although past work indicates such effects are minimal [29] and frozen slurry contains abundant, species-rich communities [15,30,31]; this is supported further by our results below.

We made and froze three replicate fresh and adapted community samples per human donor (Sup. Fig. 1). We then thawed these frozen community samples at room temperature and under anaerobic conditions. Prior to the next step, we adjusted the cell densities in the samples so they were approximately the same, estimating bacterial density by flow cytometry as described above. Then, to produce different combinations of fresh/adapted community and fresh/spent supernatant, we carried out the following steps, all in an anaerobic chamber. First, we transferred two 1.2 ml aliquots of each sample to two 2 ml tubes and centrifuged at 5000 rpm for 10 min. This gave two tubes for each sample, containing either fresh community+fresh supernatant or adapted community+spent supernatant (with three samples per human donor of both types). We used these to create all possible combinations of fresh/adapted community and fresh/spent supernatant for each human donor, in triplicate (Sup. Fig. 1). We did this by syringe filtering each supernatant either back into its original tube or into a different tube (e.g., to combine spent supernatant with fresh community), before resuspending all the pellets. We then added to each of these cultures 5 µl of overnight focal *E. coli* culture (approximately 10^7^ CFU ml^-1^), mixed by vortexing, took samples, transferred to Hungate tubes, sealed with rubber septa and incubated at 37°C static. We plated samples on chromatic agar supplemented with streptomycin to enumerate the focal strain, both at the start and after 24h.

### Amplicon sequencing

We extracted the DNA for amplicon sequencing with the powerlyzer power soil kit (Qiagen) with some modifications of the manufacturer’s protocol. In brief, we thawed samples from 0h and 72h from each microcosm of the community treatments of the main experiment and homogenized them by vortexing for 5 min. We transferred 1.5 ml of each sample into a Power Bead Tube and centrifuged at 13000 rpm for 10 min. We removed the supernatant and repeated this step to concentrate the samples. We then extracted the DNA from these concentrated samples following the manufacturers protocol. DNA quality and yield was checked with Nanodrop and Qubit.

For library preparation and sequencing, we followed the Illumina 16S Metagenomic Sequencing Library preparation guide for the MiSeq Illumina sequencing platform. We amplified the target region V3 and V4 of the 16S rRNA [32] by a limited 17 cycle PCR. We cleaned up the PCR products and attached the Nextera XT index adaptors of the Nextera XT Index Kit v2 in a second PCR step. After a clean-up of the second PCR step, we assessed quality of the library by TapeStation 2200 (Agilent) and quantified it by qPCR using the KAPA Library quantification kit on a Roche LightCycler. Libraries were then normalized, pooled and sequenced on the MiSeq sequencing platform using a 2 × 300 v3 kit, following standard Illumina sequencing protocol. We used Trimmomatic to filter raw sequencing reads and to remove adaptors. We used the scripts from the Usearch [33] software to merge amplicons into pairs, trim primer sites and clustering of OTUs. We assigned taxonomy of the Operational Taxonomic Units (OTUs) using Syntax and the SILVA 16S rRNA database (accessed April 2020). We used the phyloseq [34] package to visualize and calculate the relative abundance of taxonomic groups on the family level and alpha diversity based on total OTU abundance on the genus level.

### Statistics

To analyse variation of focal strain abundance among treatment groups in the main microcosm experiment, we used a linear mixed effects model (lmer function in R v.3.5.1 [35]). We excluded the basal medium treatment from the analysis and used Time, Donor and Community as fixed factors and Replicate as a random effect, employing a Box Cox transformation of the response variable (ƛ = 0.173439) to account for heteroscedasticity. We obtained P values for interaction terms in the model by using type II Wald chisquare test. For principal coordinate analysis of the amplicon sequencing data, we used the ordinate function of the phyloseq package with MDS as the ordination method and using Bray-Curtis dissimilarity to create the distance matrix. To test whether supernatants from different treatments varied in their effects on growth of the focal strain, we used analysis of variance (lm function) with focal strain change in CFU/ml over time as the response variable, and Donor and Treatment as factors. We ran this model both including and excluding the control treatment, testing respectively for a difference between fresh and spent medium (control vs other treatments) and between community and community-free spent-medium treatments. We used a linear mixed effects model (lmer function) for testing the effects of community age (fresh vs adapted) and supernatant age (fresh vs spent), with focal strain growth (total change in CFU count over time) as the response variable, which we log transformed and Donor, Community age and Supernatant age as fixed factors. To account for dependencies between the treatments due to the swapping of supernatant and community samples (each sample was used in two assay microcosms), we assigned a unique identifier (ID) for each independent community and supernatant sample, and used these IDs as two separate random effects. We reduced the model by removing non-significant (P>0.05) interactions using F-tests.

## Results

### *Resident microbiota suppress invasion by a focal* E. coli *strain*

We cultivated our focal *E. coli* strain in sterilized and ‘live’ versions of faecal slurry from three different human donors, measuring focal strain abundance over 72 h with periodic sampling, but without serial passage. We found the presence of resident microbial communities (in live faecal slurries) suppressed focal strain abundance compared to community-free (sterilized) versions of the same faecal slurries (effect of community in linear mixed effects model: *χ*^2^ = 41.23, d.f. = 1, *P* < 0.001; Fig. 1). This suppressive effect of resident communities was strongest toward the end of the experiment (community × time interaction: *χ*^2^ = 55.24, d.f. = 3, *p*<0.0001), with the sharpest decline between 48h and 72h, in contrast with the sterilized slurries and control treatment, where focal strain abundance was relatively stable (Fig. 1). In two community-treatment microcosms, each from a different human donor, suppression was strong enough to push the invading focal strain below our detection limit, amounting to full colonization resistance. Although average focal strain abundance varied among the different donor treatments (effect of human donor in linear mixed effect model: *χ*^2^ = 17.27, d.f. =2, *p*<0.0001), community suppression was on average consistent across human donors (community × donor interaction: *χ*^2^ = 1.64, d.f. = 2, *p*=0.44). Note the decline of focal strain abundance over time was not associated with a general decline of total bacterial population densities (including the resident microbiota): total bacterial density estimated by flow cytometry was stable at high numbers throughout the experiment (Fig. 2). This shows interactions with resident microbial communities resulted in a decrease in focal strain density over time.

**Figure 1:**
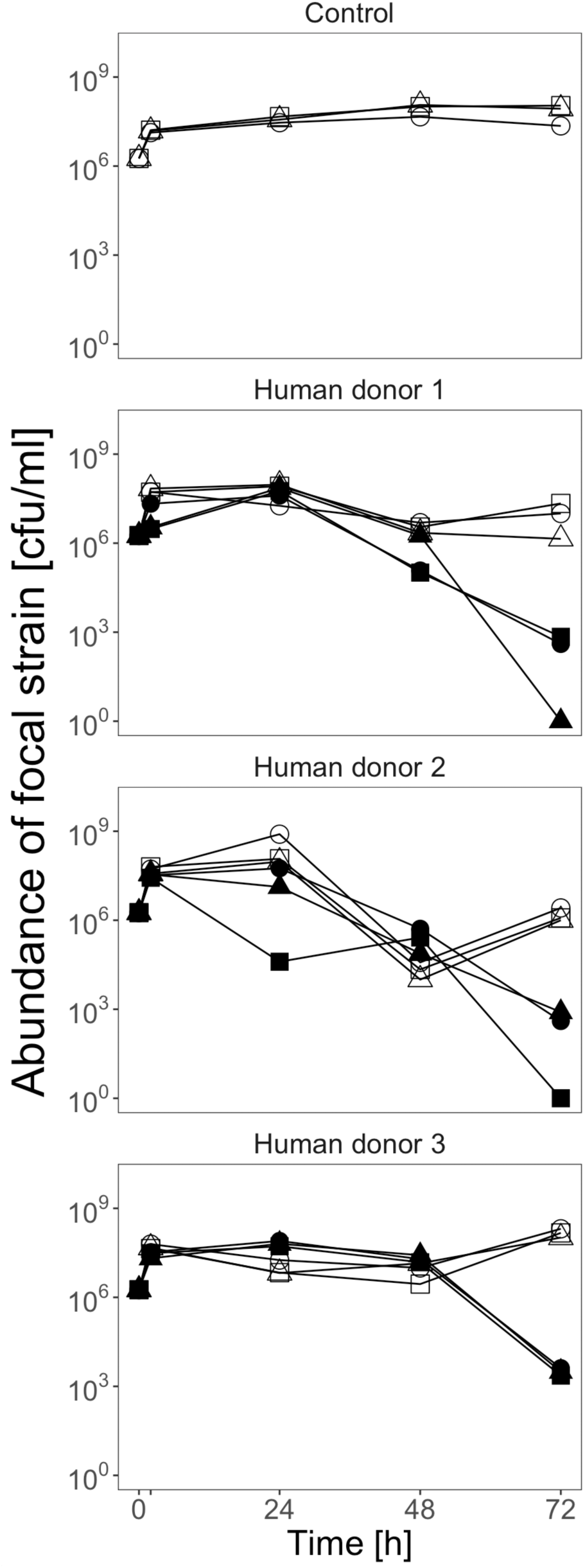
Suppressed growth of an invading focal strain in the presence of resident, human-associated microbiota. Each panel shows abundance of the focal *E. coli* strain over 72h in basal medium only (top panel) or in the presence of faecal slurry prepared with samples from one of three healthy human donors (lower three panels). For each human donor, three replicate microcosms (different symbols) are shown with sterilized faecal slurry (empty symbols) and live slurry including the resident microbiota (filled symbols). Treatments where the focal strain was below the detection limit are shown at 10°.

**Figure 2:**
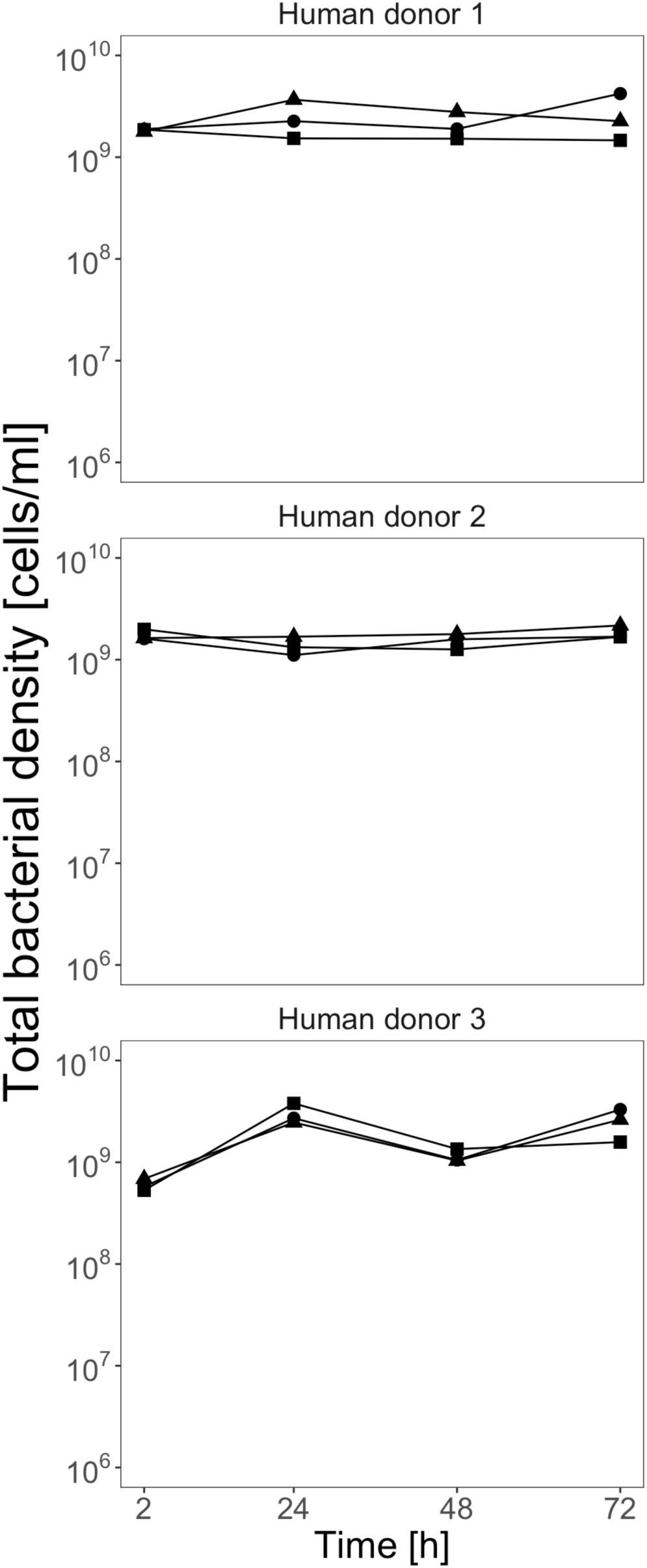
High total bacterial abundance over time. Each panel shows total bacterial abundance estimated by flow cytometry for microcosms from the community (live slurry) treatment for one of the three human donors. Replicate microcosms have different symbols.

### Resident microbial diversity was maintained over time, but with shifts in relative abundance

The stronger suppression of the invading focal strain we observed at later time points could potentially be explained by changes in the taxonomic composition of resident microbial communities. As a first step to investigate this, we tested for changes in taxonomic composition using amplicon sequencing. Taxonomic richness, taken as the number of OTUs at the genus level, of all communities was approximately stable over time (Fig. 3A). However, a measure of taxonomic diversity that accounts for evenness across different groups (Shannon index) showed a decline after 72h, and this was true for resident microbial communities from all three human donors (Fig. 3B). These shifts in relative abundance were also evident when we looked at the identities of the most abundant taxa. Initially, communities were dominated by Bacteroidaceae, Ruminococcaceae and Lachnospiraceae. Over time, these became less abundant relative to Enterobacteriaceae and Bifidobacteriaceae (Fig. 3C). Despite these changes in relative abundance, the top 10-15 families were the same after 72h. A closer look at the taxonomic assignments of the sequencing reads in the Enterobacteriaceae family revealed that almost 100% were assigned as *E. coli* and <0.1% were not annotated to the genus level. Note reads assigned to Enterobacteriaceae include those from both resident *E. coli* and the invading focal strain. To gain a rough indication of the abundance of focal strain relative to other *E. coli*, we combined information about the fraction of the total community made up by Enterobacteriaceae (from amplicon sequencing), total community abundance (from flow cytometry), and focal strain abundance (from selective plating). This suggested that in all samples the *E. coli* population was dominated by resident strains and our focal strain contributed less than 0.001% to the total *E. coli* abundance after 72h (Sup Table 1).

**Figure 3:**
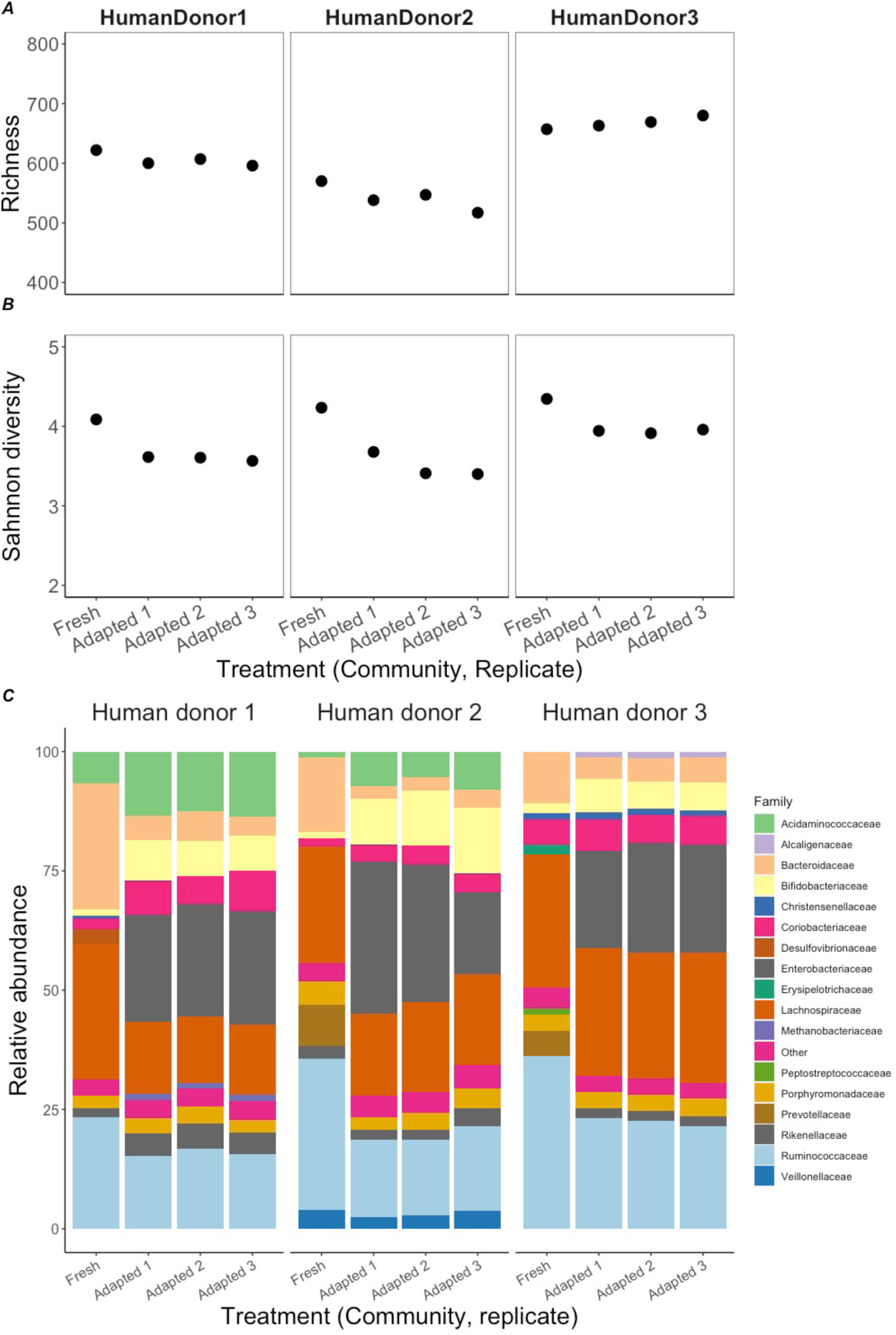
High within-sample taxonomic diversity and shifts of relative abundances over time. Taxonomic diversity based on amplicon sequencing of the 16S rRNA gene was estimated by (A) richness measured as total number of OTUs on the genus level and (B) Shannon’s diversity index at the beginning and end of the microcosm experiment. (C) Relative abundances of the top 15 families in each microcosm. For each human donor (left, middle, right panels), each of these measures is shown for the fresh faecal slurry (start of the microcosm experiment) and three replicate ‘adapted’ microcosms from the community treatments at the end of the microcosm experiment (*x*-axis).

Principal coordinate analysis (PCoA) based on Bray-Curtis dissimilarities confirmed that resident microbial communities in different microcosms changed over time in similar ways (along the same axis in Sup. Fig. 2A). This also revealed that communities from human donors 1 and 2 were more similar to each other than to those from human donor 3 (Sup. Fig. 2A). An alternative PCoA analysis excluding Enterobacteriaceae from the dataset showed a similar qualitative trend, suggesting the expansion of this family alone does not explain the shift in community composition (Sup. Fig. 2B). In summary, our analysis of total microbial diversity revealed shifts in relative abundance for certain taxa, such as an expansion of resident *E. coli* strains, and these coincided with increased suppression of the focal strain toward the end of the experiment observed above.

### Changing abiotic conditions alone do not explain focal strain suppression

Another possible explanation for the observed suppression of the invading focal strain, which does not require changes in community composition, is that the resident microbiota change the local abiotic environment over time in a way that permits less focal strain growth than in the community-free (focal strain only) treatments (e.g., resource depletion or accumulation of compounds toxic to the focal strain). To test this, we inoculated our focal strain into supernatant extracted by centrifugation and filtration of cultures from the community and community-free treatments at the end of the main microcosm experiment above, and into freshly prepared basal medium supplemented with sterilized slurry (unspent medium with slurry thawed from frozen, equivalent to the medium used in community-free treatments at the start of the main experiment) as a control. We found our focal strain grew in the supernatants from both the community- and community-free treatments (positive net change in abundance over 24h; Fig. 4). This shows that in neither treatment had the environment become toxic to the focal strain. Furthermore, supernatants from both treatments supported a similar amount of growth of the focal strain (effect of Treatment in a model excluding the control treatment: F_1,12_=1.4, *p*=0.26). This shows mixed communities comprising the resident microbial community plus the focal strain did not deplete available nutrients that support focal strain growth any more than the focal strain did when growing alone. Despite this, the supernatants from both of these treatments supported significantly less growth compared to freshly prepared, sterilized-faecal-slurry medium (effect of Treatment, including the control treatment: F_2,18_=28.7, *p*< 0001). This is consistent with microbial activity in our main microcosm experiment depleting resources that the focal strain uses for growth. We refer to this hereafter as nutrient depletion (for the focal strain) but, as we discuss further below, this does not necessarily mean the environment became nutrient poor for all resident bacteria (not the focal strain) as well. Moreover, this does not explain why we observed suppression of the focal strain in the community treatments compared to the community-free treatments.

**Figure 4:**
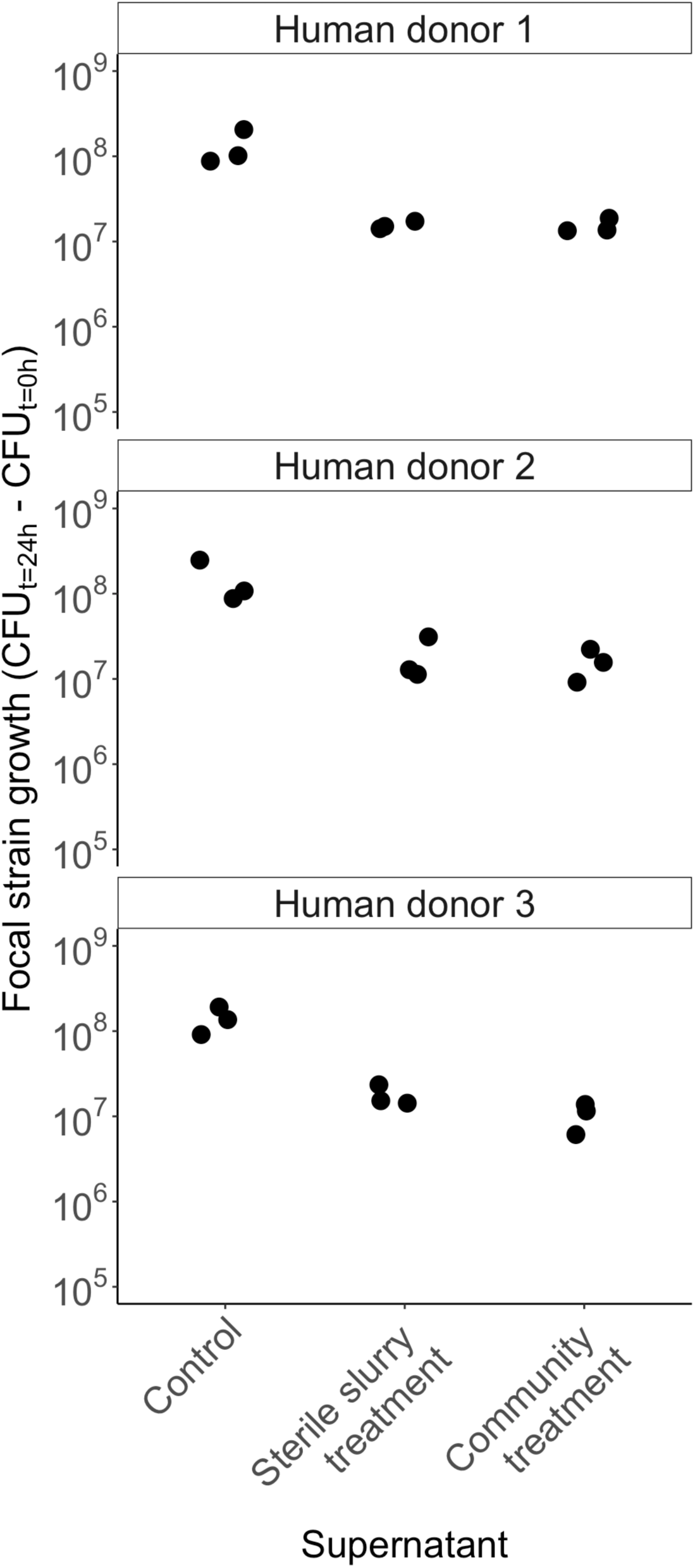
Nutrient depletion in both community and community-free treatments. Focal strain growth (change in CFU density over time) in freshly prepared supernatant (equivalent to community-free treatment at the start of the main experiment, Fig. 1), or supernatant from the community and community-free treatments at the end of the main experiment. The three replicates in each treatment had supernatant from independent microcosms of the main experiment.

### Suppressing effects of resident communities depend on local abiotic conditions

Having observed changes in both community composition and the local abiotic environment (nutrient depletion) above, we hypothesised that suppression of the invading focal strain toward the end of the main experiment resulted from an interaction between these two factors. That is, we asked whether suppression required communities that had adapted to our microcosm environment, depleted nutrient status in the microcosm, or both. To do this, we compared fresh and adapted versions of the same resident microbial communities used above (prepared from frozen faecal slurry, and sampled before and after 72h incubation in the same conditions as in the main experiment, but without the focal strain) and fresh and spent versions of the local abiotic environment (by extracting supernatant from the same microcosms used to prepare fresh and adapted communities). We then made a fully factorial experiment testing the effects of fresh/adapted communities and fresh/spent supernatant on focal strain growth (Sup. Fig. 1). This showed adapted communities suppressed focal strain growth compared to fresh communities on average, but only in spent medium (community × medium interaction: *χ*^2^ = 179.57, d.f. = 1, *p*<0.001; Fig. 5). This is in line with the main microcosm experiment, where the human donor 2 community showed stronger suppression of the focal strain already after 24h compared with the communities from human donors 1 and 3. In summary, changes in community composition over time made adapted microbiota more suppressive than fresh microbiota, but for two out of three human donors, this was only observed after the abiotic conditions had become relatively nutrient poor for the focal strain.

**Figure 5:**
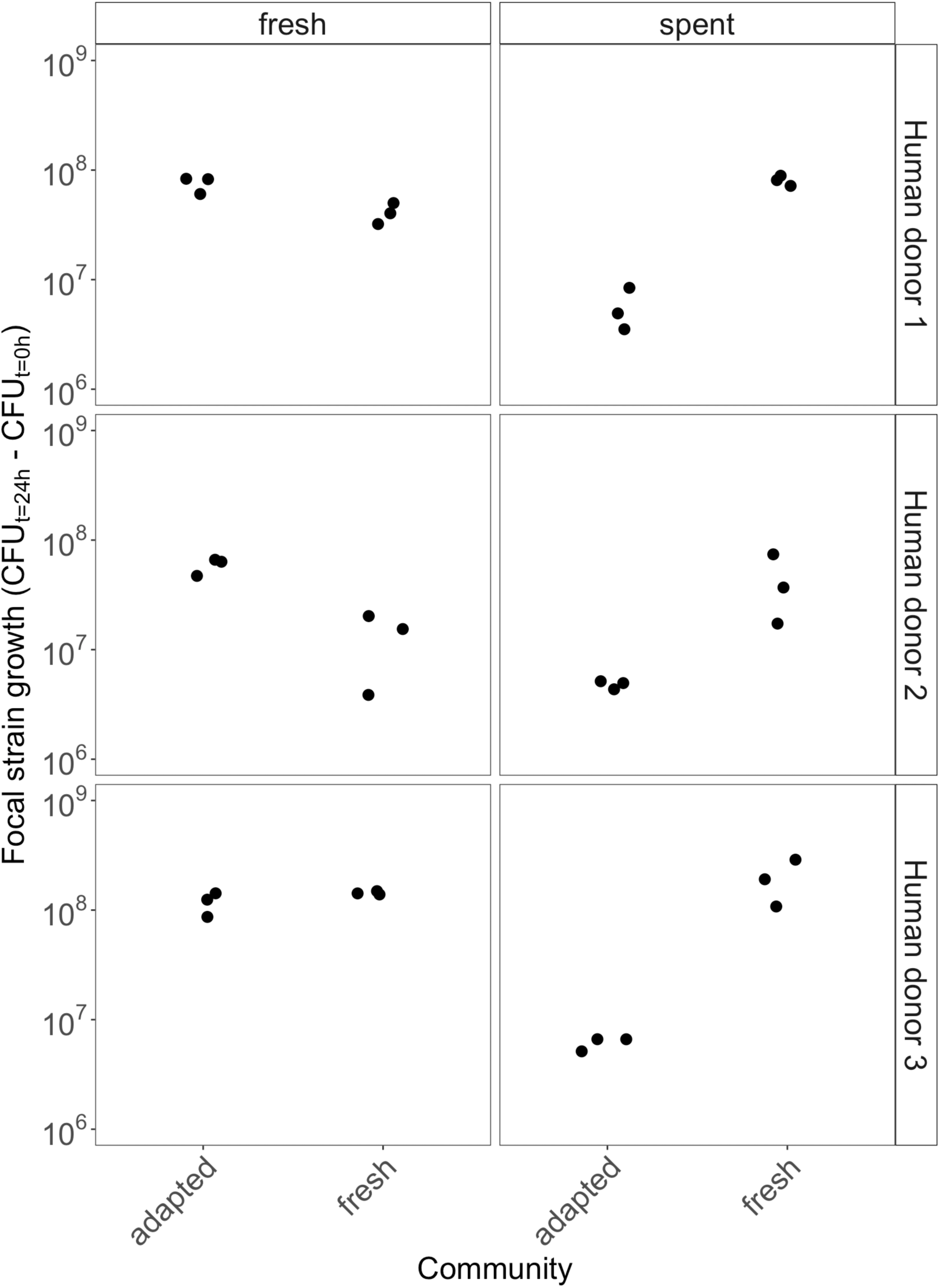
Focal strain suppression depends on community composition and the state of the environment. Each panel shows growth over 24h of the focal *E. coli* strain (in CFU ml^-1^) co-cultivated either with an adapted community or with a community coming directly from faecal slurry from each human donor. Each co-culture was either inoculated into fresh medium (left column) or spent supernatant from 72h-old microcosms (right column).

## Discussion

We showed colonization success of an invading lineage depended on an interaction between the taxonomic composition of the resident community and the nutrient status of the local environment. We demonstrated this by monitoring invasion success of a strain of the common gastrointestinal species *E. coli* in microbiome samples from humans, cultivated in anaerobic microcosms. By comparing live and deactivated versions of the same samples, we showed time-dependent suppression of the invading *E. coli* strain, in some cases amounting to full colonization resistance (extinction of the invader). This suppression coincided with a shift in microbial community composition and declining nutrient status (for the invading focal strain) in the microcosms. However, by splitting the microcosm system into the liquid phase and the resident microbes (by centrifugation and filtration), we showed these two factors interact with each other. Microbial communities that had adapted to their local microcosm environments were most resistant to invasion, particularly when available resources were scarce, amplifying competition with invading strains. This provides new insights into what makes some communities more susceptible to invasion than others.

The first key implication of our results is the dependency of colonization resistance on both aspects, community composition and local abiotic conditions. That changing taxonomic composition was linked to changes in susceptibility to invasion is promising in terms of predicting colonization resistance of microbiota from individual people. However, translating taxonomic information such as 16S data into predictions about community-level susceptibility to invasion by pathogens remains a significant challenge [36]. Summary metrics, such as species diversity as we analyzed above, may help here, and have been correlated with some microbiota-related properties such as risk of obesity [37], diabetes [38] and recurrent infection with *Clostridium difficile* [39]. However, in our experiment diversity by some measures (Shannon index) declined as communities became more robust to invasion, and by others (richness) remained stable. This is in apparent contrast with the general principle that more diverse ecosystems should be harder to invade [2,40]. Thus, it is probably not simply the case that higher diversity means more colonization resistance. In support, previous studies in mouse models identified consortia of four and 15 commensal species that cleared the intestine from an antibiotic-resistant *Enterococcus faecium* strain and established colonization resistance against *Salmonella enterica* serovar Typhimurium [41,42]. More importantly, our finding that suppressive effects of resident communities were strongly modified by local conditions (different supernatants), suggests sequence data alone, or other information about community composition, are insufficient to accurately predict colonization resistance. This is consistent with past work showing nutrient supplementation can interfere with competitive interactions among resident and invading microbes in mouse microbiota [43,44]. Our results go beyond this to show nutrient status also plays a key role in modulating colonization resistance in human-associated microbiota.

The second key implication of our work is for interventions aimed at improving colonization resistance of individual hosts/patients, such as faecal microbial transplantation [10]. The interaction between community composition and local abiotic conditions we observed indicates colonization resistance resulting from such interventions will depend not only on the type of community that is implanted, but on factors that influence the local micro-environment, such as host diet or physiological status. We found this interaction varied among donors (with donor 2 community samples being relatively suppressive even with fresh supernatant), indicating some person-to-person variation of the relative importance of different drivers of colonization resistance. However, that suppression of the invading focal strain was consistent across donors is also encouraging, indicating some key properties are repeatable across randomly selected healthy-donor communities. This, and our amplicon data, are consistent with the notion of a core microbiome conferring similar functions across healthy individuals, despite variation of the individual taxa present [45].

Part of the taxonomic shift over time in our main experiment was driven by expansion of Enterobacteriaceae, the same family our focal strain belongs to. Therefore, intraspecific competition between resident and focal *E. coli* may explain at least some of the suppressive effects of resident communities. Consistent with this, we found previously that resident *E. coli* strains sampled from other human donors had a competitive advantage over this focal strain *in vitro* [14]. Turnover of resident and transient *E. coli* clones has recently been observed in samples from humans [46], suggesting such intraspecific competition is also possible in nature. Nevertheless, other taxa also increased in relative abundance, such as Bifidobacteriaceae (Fig. 3), so we do not exclude there also being a role for interspecific competition. More importantly, the ecological mechanism behind our key results (stronger suppression of an invading strain when the resources it uses for population growth are scarce and the resident community is adapted to local conditions), is likely not limited to particular strains or species. Other *E. coli* strains may be stronger or weaker invaders compared to the focal strain we used, but crucially this set-up enabled us to observe variable invasion success depending on interactions with resident microbiota and local abiotic conditions.

Using replicated microcosms with live and inactivated faecal slurry allowed us to directly observe the effects of human-associated microbiota on an invading focal strain. This set-up also imposes some important limitations. First, we account here only for drivers of colonization resistance involving microbial interactions, not those that also involve the host immune system [13], which may influence the composition of the gut microbiota. Despite this, our findings are consistent with evidence that competition in nutrient-depleted conditions also matters for colonization resistance in mice [47–49], although not always [50]. A second limitation of our approach is that our centrifuged and filtered supernatants, although clearly containing far less bacteria than the pelleted part of each sample, could potentially have contained other bioactive material such as bacteriophages. While we do not exclude the possibility that bacteriophages in our supernatants could have infected the focal strain in subsequent assays, this seems unlikely to explain our results. Such bacteriophages would not have been present in supernatants from the sterile slurry treatments (Fig. 4), where we observed similar suppression compared to supernatants from community treatments. We therefore found no evidence that infectious material in community-treatment supernatants altered their effect on focal strain growth compared to nutrient depletion alone. Additionally, in a previous study we screened faecal samples collected using a similar design for plaque-forming units with this focal strain, and found none [14].

In summary, our results suggest the outcome of invasion by a new strain or species depends strongly on the taxonomic composition of the resident human gut microbiota. 16S rRNA amplicon analysis revealed changes in community composition over time that coincided with nutrient depletion in the local abiotic environment and stronger suppression of the invading lineage. However, nutrient depletion alone did not explain suppression of the invading strain. Instead, altered abiotic conditions contributed to competitive suppression only when the resident community had also adapted to the conditions of our experiment. A key challenge for putting such insights into practice in the context of microbiota-based treatments is to identify scenarios (types of infections, health conditions or other biomarkers) where susceptibility to infection can be predicted more strongly from taxonomic information, and interventions that can change the within-host environment in ways that maximise colonization resistance against pathogens.

### Data availability

16S rRNA sequences are deposited in the European Nucleotide Archive under the accession number PRJEB38488. Raw data of the experiments will be available through the Dryad repository upon publication.

## Acknowledgements

We thank the Genetic Diversity Center and Jean-Claude Walser for sequencing and bioinformatic support, and Pauline Scanlan for helpful feedback. Funded by the Swiss National Science Foundation project 31003A_165803.

## Figure legends

**Sup. Fig. 1:**
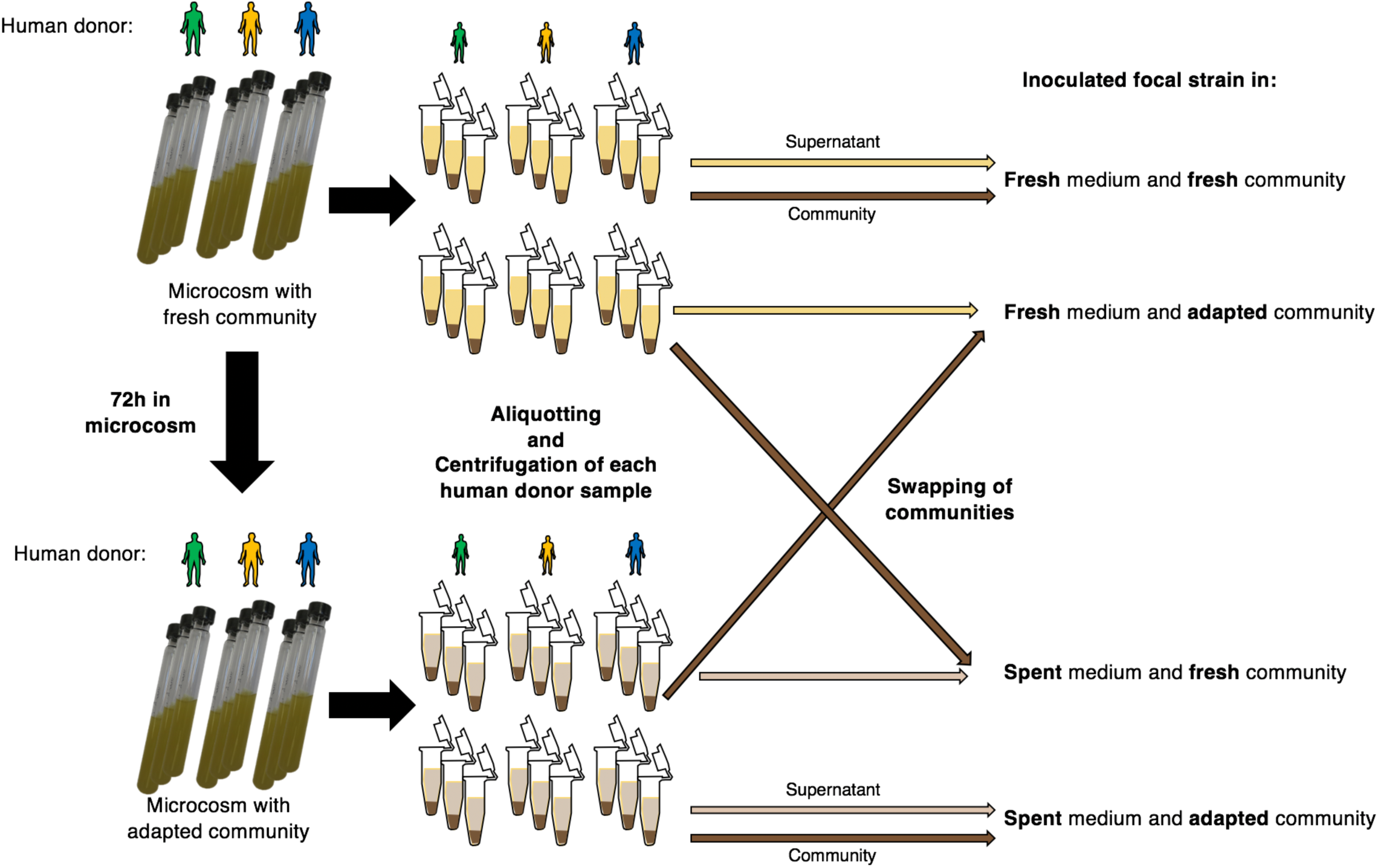
Schematic of experimental swapping of communities and supernatants. To produce fresh/adapted community samples and fresh/spent medium, fresh faecal slurry from the three different human donors was inoculated into freshly prepared basal medium, as at the start of the main experiment (Fig. 1), but without the focal strain. Fresh community/fresh medium samples were frozen immediately; adapted community/spent medium samples were frozen after 72h incubation in the same conditions as in the main experiment. We made three replicate microcosms of each type. To separate communities and media (supernatants), we thawed the samples and took aliquots of each, before centrifuging them. Then, in a fully factorial design (all combinations of fresh/adapted community and fresh/spent medium within each donor and replicate), we resuspended the supernatant from each centrifuged sample, either into the same tube or into another tube. We then inoculated these samples with the focal strain (see Methods) and incubated for 24 h under static conditions at 37 °C. This was all done under anaerobic conditions.

**Sup. Fig. 2:**
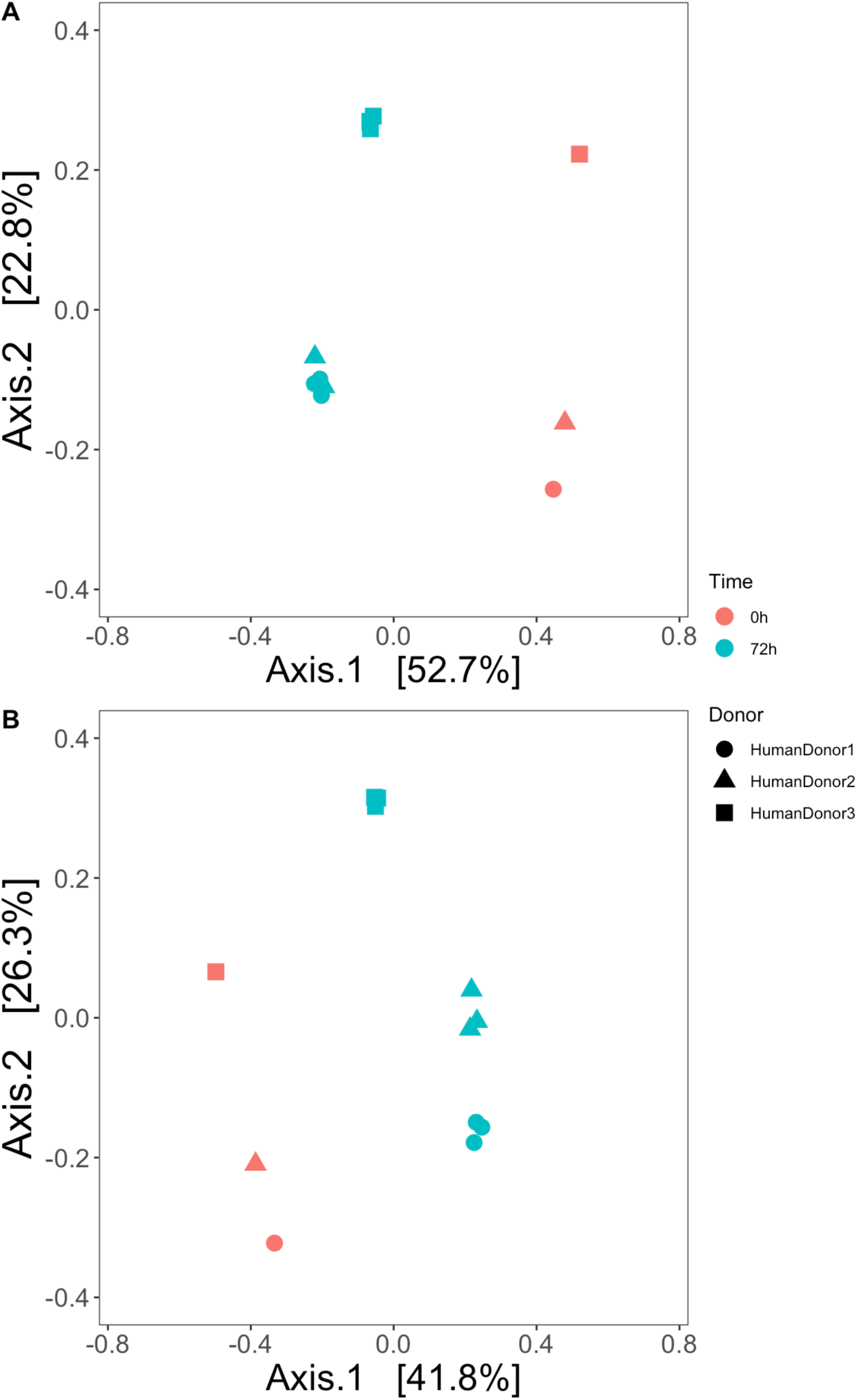
Changes in microbial composition over time and across human donors. **(A)** This panel shows ordination of the entire amplicon sequence dataset, with samples from each initial community (red symbols) and samples taken after 72h (blue symbols); samples from the three human donors are shown with different symbol shapes. **(B)** The same ordination but excluding Enterobacteriaceae. In both panels, Bray-Curtis distance was used for the distance matrix to calculate similarities and Principal coordinate analysis to plot the results.

## Sup. Table 1

**Sup Table 1:**
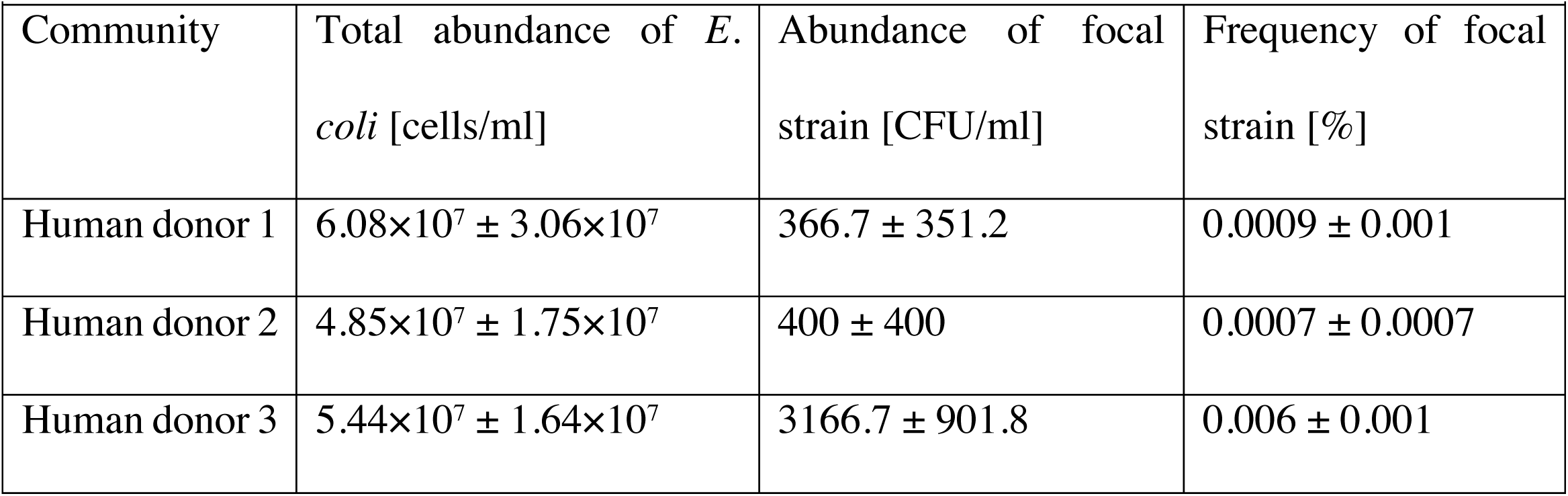
Abundance of total Enterobacteriaceae and focal strain at the end of the 72h microcosm experiment. *Community* gives the origin of the microbiome sample. *Total abundance of E. coli* was estimated from total bacterial abundance measured by flow cytometry and the frequency of total *E. coli* based on the reads distribution of the amplicon data. *Abundance of focal strain* was estimated by selective plating. *Frequency of focal strain* is the ratio of the focal strain divided by the total abundance of *E. coli*. Each value is the mean ±s.d. from three replicate microcosms. Note we interpret these estimates from combining data types with caution, and take them only as a rough indication of whether *E. coli* was rare or common relative to other *E. coli*.

